# Achieving stable dynamics in neural circuits

**DOI:** 10.1101/2020.01.17.910174

**Authors:** Leo Kozachkov, Mikael Lundqvist, Jean-Jacques Slotine, Earl K. Miller

## Abstract

The brain consists of many interconnected networks with time-varying, partially autonomous activity. There are multiple sources of noise and variation yet activity has to eventually converge to a stable, reproducible state (or sequence of states) for its computations to make sense. We approached this problem from a control-theory perspective by applying contraction analysis to recurrent neural networks. This allowed us to find mechanisms for achieving stability in multiple connected networks with biologically realistic dynamics, including synaptic plasticity and time-varying inputs. These mechanisms included inhibitory Hebbian plasticity, excitatory anti-Hebbian plasticity, synaptic sparsity and excitatory-inhibitory balance. Our findings shed light on how stable computations might be achieved despite biological complexity.

## 2 Introduction

Behavior emerges from complex neural dynamics unfolding over time in multi-area brain networks. Even in tightly controlled experimental settings, these neural dynamics often vary between identical trials^1,2^. This can be due to a variety of factors including variability in membrane potentials, inputs, plastic changes due to recent experience and so on. Yet, in spite of these fluctuations, brain networks must achieve computational stability: despite being “knocked around” by plasticity and noise, the behavioral output of the brain on two experimentally identical trials needs to be similar. How is this stability achieved?

Stability has played a central role in computational neuroscience since the 1980’s, with the advent of models of associative memory that stored neural activation patterns as stable point attractors^3–7^, although researchers were thinking about the brain’s stability since as early as the 1950’s^8^. The vast majority of this work is concerned with the stability of activity around points, lines, or planes in neural state space^9,10^. However, recent neurophysiological studies have revealed that in many cases, single-trial neural activity is highly dynamic, and therefore potentially inconsistent with a static attractor viewpoint^1,11^. Consequently, there has been a number of recent studies—both computational and experimental—which focus more broadly on the stability of neural *trajectories*^12,13^.

While these studies provide important empirical results and intuitions, they do not offer analytical insight into mechanisms for achieving stable trajectories in recurrent neural networks. Nor do they offer insights into achieving such stability in plastic (or multi-modal) networks. Here we focus on finding conditions that guarantee stable trajectories in recurrent neural networks and thus shed light onto how stable trajectories might be achieved *in vivo*.

To do so, we used contraction analysis, a concept developed in control theory^14^. Unlike a chaotic system where perturbations and distortions can be amplified over time, the population activity of a contracting network will converge towards the same trajectory, thus achieving stable dynamics (Figure 1). One way to understand contraction is to represent the state of a network at a given time as a point in the network’s ‘state-space’, for instance the space spanned by the possible firing rates of all the networks’ neurons. This state-space has the same number of dimensions as the number of units *n* in the network. A particular pattern of neural firing rates corresponds to a point in this state-space. This point moves in the *n* dimensions as the firing rates change and traces out a trajectory over time.

**Figure 1:**
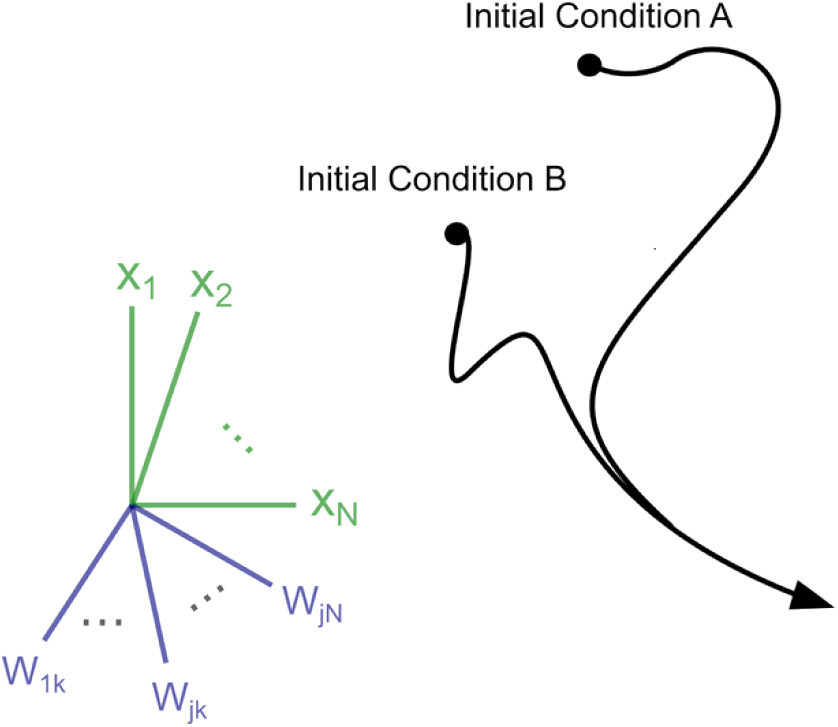
Cartoon demonstrating the contraction property. In a network with *N* neural units and *S* dynamic synaptic weights, the network activity can be described a trajectory over time in an (*N* + *S*)-dimensional space. In a contracting system all such trajectories will converge exponentially *in some metric* towards each other over time, regardless of initial conditions. In other words, the distance between any two trajectories shrinks to **zero—**potentially after transient divergence (as shown).

In a contracting network, all such trajectories converge. These contracting dynamics have previously been used in several applications, including neural networks with winner take all dynamics^15,16^, in a model of action-selection in the basal ganglia^17^, and to explain how neural synchronization can protect from noise^18^. Here, we instead explore how contraction can be achieved generally in more complex recurrent neural networks (RNNs) including those with plastic weights. We used RNNs that received arbitrary time-varying inputs and had synapses that changed on biologically relevant timescales^19–21^. Our analysis reveals several novel classes of mechanisms that produced contraction including inhibitory Hebbian plasticity, excitatory anti-Hebbian plasticity, excitatory-inhibitory balance, and sparse connectivity.

## 3 Results

We used two main quantitative tools to characterize contraction. One was the contraction *rate*. It indicates how fast trajectories reconvened following a perturbation. Another was the Jacobian of the networks. The Jacobian of a dynamical system is a matrix essentially describing the local ‘traffic laws’ of nearby trajectories of the system in its state space. More formally, it is the matrix of partial derivatives describing how a change in any system variable impacts the *rate of change* of every other variable in the system. It was shown in^14^ that if the matrix measure—also known as the logarithmic norm^22^ – of the Jacobian is negative then all nearby trajectories are funneled towards one another (see A.1.2 for technical review). This, in turn, implies that *all* trajectories are funneled towards one another. The contraction rate, how quickly a perturbation is forgotten, and the matrix measure are related as follows: The maximum value attained by the matrix measure of the Jacobian along the network’s trajectory *is* the contraction rate.

Importantly, the above description can be generalized to different *metrics*. In other words, if one can find a (potentially time-varying) coordinate system in which the network is contracting—in the sense that its Jacobian *in the new coordinates* is negative definite-this implies contraction for *all* coordinate systems. This makes contraction analysis useful for analyzing systems where exponential convergence of trajectories is preceded by transient divergence (Figure 1) as in recent models of motor cortex^23,24^. In this case, it is usually possible to find a coordinate system in which the convergence of trajectories is ‘pure’. For example, linear stable systems were recently used in the motor control literature to find initial conditions which produce the most energetic neural response^23^ They are ‘purely’ contracting in a metric defined by the eigenvectors of the weight matrix (see Example 5.1 in^14^) but transiently diverging in the identity metric.

### 3.1 Inhibitory Hebbian Plasticity & Excitatory Anti-Hebbian Plasticity Produce Contraction

It is known that certain forms of synaptic plasticity can quickly lead to extreme instabilities if left unchecked^9,25^. Thus, the same feature that can aid learning can also yield chaotic neural dynamics if not regulated. It is not known how the brain resolves this dilemma. A growing body of evidence—both experimental and computational—suggests that inhibitory plasticity (that is, the strengthening of inhibitory synapses) can stabilize neural dynamics while simultaneously allowing for learning/training in neural circuits^26–28^. By using the Jacobian analysis outlined above, we found that inhibitory Hebbian synaptic plasticity (as well as excitatory anti-Hebbian plasticity) indeed leads to stable dynamics in neural circuits. Specifically, we considered neural networks of the following form:

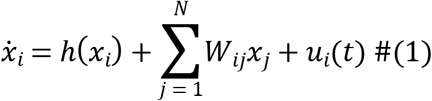

The term 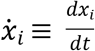 denotes the change in the activation of neuron *i* as a function of time and *W_ij_* denotes the weight between neurons *i* and *j*. The term *h*(*x_i_*) captures the ‘self-dynamics’ of neuron *i* —the dynamics it would have in the absence of input from other neurons. The term being summed represents the weighted contribution of all the neurons in the network on the activity of neuron *i*. Finally, the term *u_i_*(*t*) represents external input into neuron *i*.

We did not constrain the inputs into the RNN (except that they were not infinite) and we did not specify the particular form of *h*(*x_i_*) except that it should be a leak term (i.e has a negative derivative for all *x*, see A.2.2.4). Furthermore, we made no assumptions regarding the relative timescales of synaptic and neural activity. Synaptic dynamics were treated on an equal footing as neural dynamics. We considered synaptic plasticity of the following form:

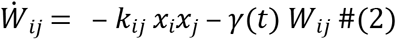

where the term *k_ij_* > 0 is the learning rate for each synapse and *γ*(*t*) > 0 is a decay factor (the rate of forgetting) for each synapse. For technical reasons outlined in the appendix (A.3.2), we restricted **K**, the matrix containing the learning rates *k_ij_*, to be positive-semidefinite, symmetric, and have positive entries. A particular example of **K** satisfying these constraints is to have the learning rates of all synapses to be equal (i.e. *k_ij_* = *k* > 0). For excitatory synapses, this equation describes anti-Hebbian plasticity because the synaptic weight becomes less positive (and thus weaker) between correlated neuron pairs. For inhibitory synapses, the above equation describes Hebbian plasticity because the direction of synaptic weight change is negative between correlated neurons, and thus the amplitude of the synapse *increase*^29,30^. Plasticity of this form produced contracting neural and synaptic dynamics regardless of the initial values of the weights and neural activity (Figure 2 and Figure 3). The red trace of Figure 3A shows that this is not simply due to the weights decaying to 0. Thus, this plasticity is not only contraction preserving, it is contracting *ensuring*. Furthermore, we showed that the network is contracting in a non-identity metric (which we derive from the system parameters in **K**), opening up the possibility of transient divergent dynamics in the identity metric.

**Figure 2:**
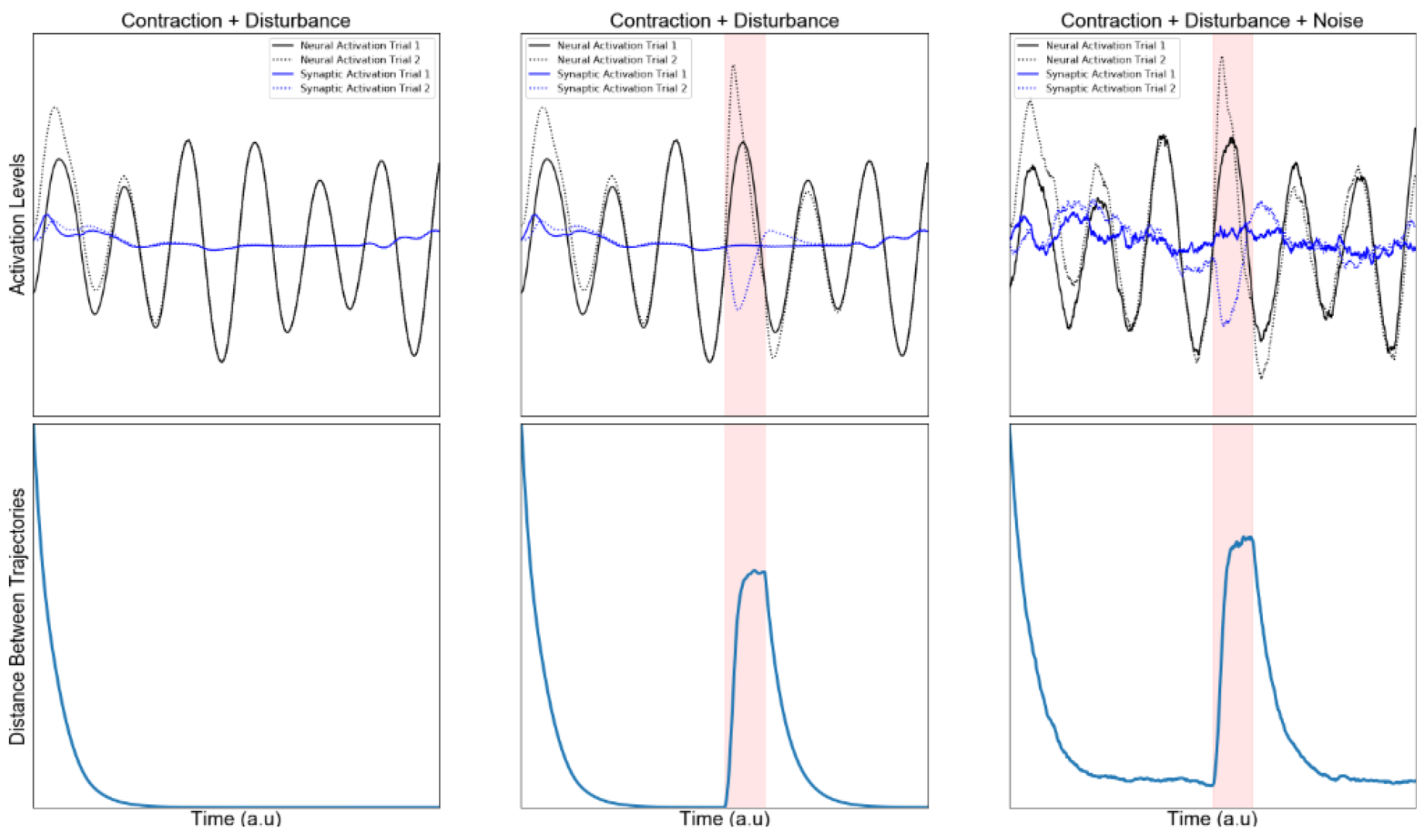
Contracting dynamics of neural and synaptic activity. Euclidean distances between synaptic and neural trajectories demonstrate exponential shrinkage over time. The top row of panels shows the activation of a randomly selected neural unit (black) and synapse (blue) across two simulations (dotted and solid line). The bottom row shows the average Euclidean distance in state space for the whole population across simulations with distinct, randomized starting conditions. Leftmost Panel: Simulations of a contracting system where only starting conditions differ over simulations. Center Panel: the same as in Leftmost but with an additional random pulse perturbation in one of the two simulations indicated by a red background shading. Rightmost Panel: same as in Center Panel but with additional sustained noise, unique to each simulation.

**Figure 3:**
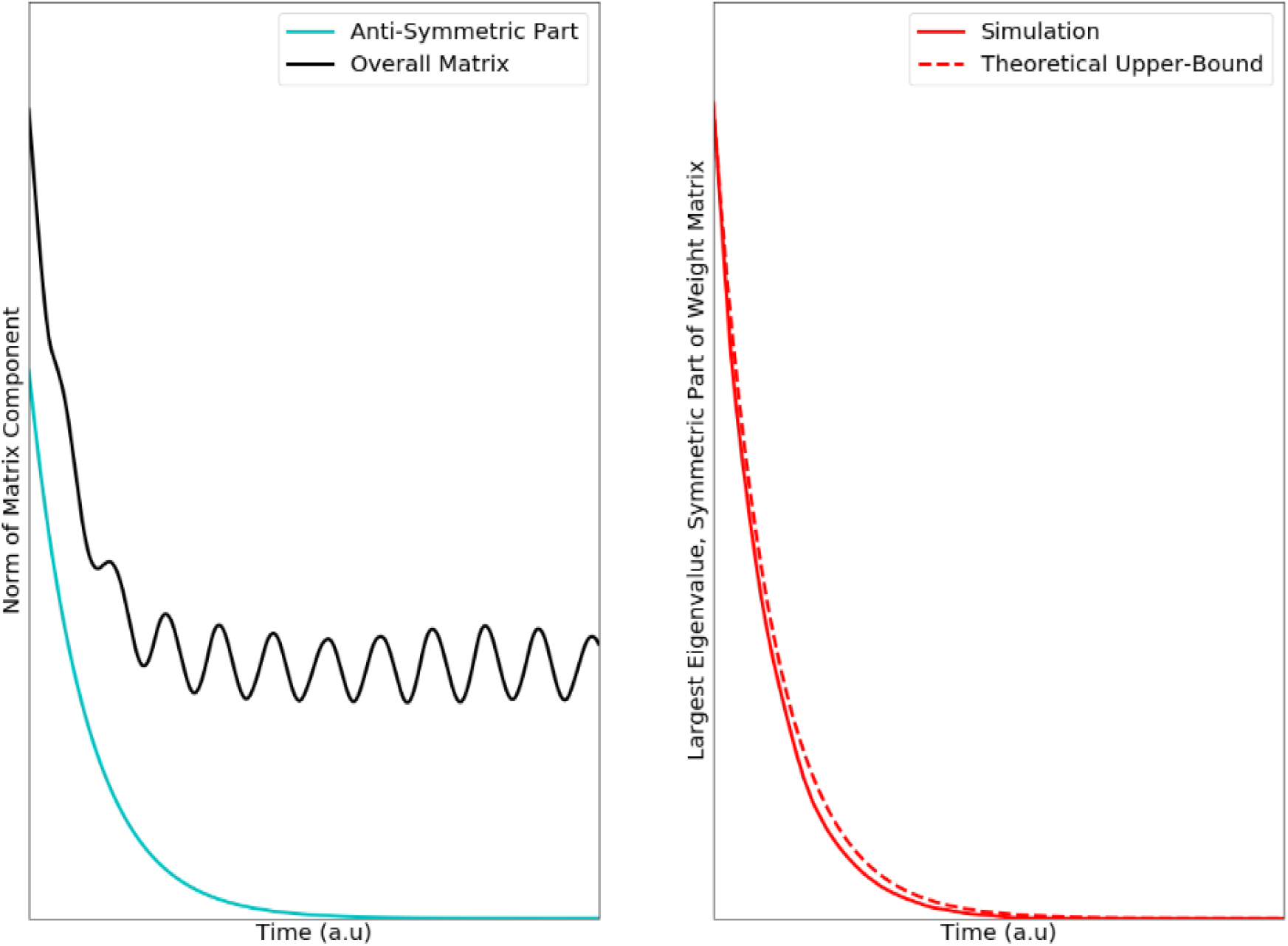
(Left) The anti-Hebbian plasticity pushes the weight matrix towards symmetry. Left, teal trace) Plotted is a measure (the spectral norm) of how asymmetric the weight matrix is. Teal curve shows that this measure decays to zero, implying the weight matrix becomes symmetric. The black trace shows spectral norm of the weight matrix. If this quantity does not decay to zero, it implies that not all the weights have decayed to zero. On the right, we plot the largest eigenvalue of the symmetric part of W. A prerequisite for overall contraction in the network is that this quantity be less than or equal to the ‘leak-rate’ of the individual neurons. The dotted line shows our theoretical upper bound for this quantity, and the solid line shows the actual value of taken from a simulation (see methods).

To explain how inhibitory Hebbian plasticity and excitatory anti-Hebbian plasticity work to produce contraction across a whole network, we needed to deal with the network in a holistic fashion, not by analyzing the dynamics of single neurons. To do so, we conceptualized RNNs with dynamic synapses as a single system formed by combining two subsystems, a neural subsystem and a synaptic subsystem. We showed that the above plasticity rule led the neural and synaptic subsystems to be independently contracting. Thus contraction analysis of the overall system then boiled down to examining the interactions between these subsystems^31^.

We found that this plasticity works like an interface between these systems. It produces two distinct effects that push networks toward contraction. First, it makes the synaptic weight matrix symmetric (Figure 3A, red trace). This means that the weight between neuron *i* to *j* is the same as *j* to *i*. We showed this by using the fact that every matrix can be written as the sum of a purely symmetric matrix and a purely anti-symmetric matrix. An anti-symmetric matrix is one where the *ij* element is the negative of the *ji* element (*i.e*. *W_ij_* = −*W_ij_*) and all the diagonal elements are zero. We then showed that anti-Hebbian plasticity shrinks the anti-symmetric part of the weight matrix to zero, implying that the weight matrix becomes symmetric. The symmetry of the weight matrix ‘cancels out’ off-diagonals in the Jacobian matrix (see A.3) of the overall neural-synaptic system. Loosely speaking, off-diagonal terms in the Jacobian represent potentially destabilizing cross-talk between the two subsystems. Furthermore, anti-Hebbian plasticity makes the weight matrix negative semi-definite. This means that all its eigenvalues are less than or equal to zero (Figure 3).

### 3.2 Sparse Connectivity Pushes Networks toward Contraction

Synaptic connectivity in the brain is extraordinarily sparse. The adult human brain contains at least 10^11^ neurons yet each neuron forms and receives on average only 10^3^-10^4^ synaptic connections^32^. That is about 18 orders of magnitude more sparse than if the brain’s neurons were all to all connected 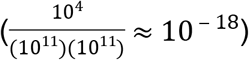. Even in local patches of cortex, such as we model here, connectivity is far from all-to-all; cortical circuits are sparse^33^. Our analyses revealed that sparse connectivity helps produce global network contraction for many types of synaptic plasticity.

To account for the possibility that some synapses may have much slower plasticity than others (and can thus be treated as synapses with fixed amplitude), we made a distinction between the total number of synapses and the total number of *plastic* synapses. These plastic synapses then changed on a similar time-scale as the neural firing rates. By neural dynamics, we mean the change in neural activity as a function of time. We analyzed RNNs with the structure:

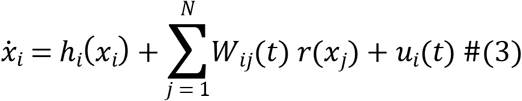

Where *h_i_*(*x_i_*) is a nonlinear leak term (see A.2.2.4), and *r*(*x_j_*) is a nonlinear activation function. The RNNs analyzed in this section are identical to those analyzed in the previous section, with the exception of the activation term. Here, we allow for a more general class of activations, whereas in the previous section we constrained *r*(*x_j_*) to be linear for analytical tractability. We denote the total number of afferent synapses into neuron *i* by *p_i_* and the number of afferent *plastic* synapses by *d_i_*. Since the number of dynamic synapses cannot be greater than the total number of synapses, *d_i_* has to be a fraction of *p_i_*, This means we can write it as *d_i_* = *α_i_p_i_*, where *α_i_* is a number between 0 and 1. We refer to the maximum possible absolute strength of a synapse as *w_max_*, the maximum possible firing rate of a neuron as *r_max_* and finally the contraction rate of the *i^th^* isolated neuron as *β_i_*. Recall from the introduction that the contraction rate measures how quickly the trajectories of a contracting system reconvene after perturbation. Under the assumption that the plastic synapses are contracting, we show in the appendix (Section 4) that if the following equation is satisfied for every neuron, then the overall network is contracting:

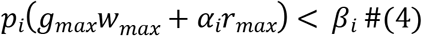

Where *g_max_* is the maximum gain of any neuron in the network (see A.4). Because *β_i_* is a positive number, it is always possible to decrease *p_i_* to the point where this equation is satisfied. Because increasing the sparsity of a network corresponds to decreasing *p_i_*, we may conclude that increasing the sparsity of connections pushes the system in the direction of contraction. This equation also implies that the faster the individual neurons are contracting (i.e. the larger *β* is), the denser you can connect them with other neurons while still preserving overall contraction.

### 3.3 E-I Balance Leads to Contraction in Static RNNs

Apart from making connections sparse, one way to ensure contraction is to make synaptic weights small. This can be seen for the case with static synapses by setting *α_i_* = 0 in the section above, where *W*_max_ now have to be small to ensure contraction. Intuitively, this is because very small weights mean that neurons cannot exert much influence on one another. If the neurons are stable before interconnection, they will remain so. Since strong synaptic weights are commonly observed in the brain, we were more interested in studying when contraction can arise irrespective of weight amplitude. Negative and positive synaptic currents are approximately balanced in biology^34–36^. We reasoned that such balance might allow much larger weight amplitudes while still preserving contraction since most of the impact of such synapses cancel and the net effect small. This was indeed the case.

To show this, we studied the same RNN as in the section above, while assuming additionally that the weights are static. In particular, we show in the appendix (Section 5) that contraction can be assessed by studying the eigenvalues of the *symmetric* part of **W** 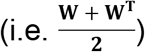. This implies the following: if excitatory to inhibitory connections are of equal amplitude (and opposite sign) as inhibitory to excitatory connections, they will not interfere with stability—regardless of amplitude (see A.5). This is because connections between inhibitory and excitatory units will be in the off-diagonal of the overall weight matrix and get cancelled out when computing the symmetric part. As an intuitive example, consider a two-neuron circuit made of one excitatory neuron and one inhibitory neuron connected recurrently (as in^37^, Fig 1A). Assume that the overall weight matrix has the following structure:

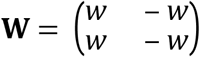

When taking that symmetric part of this matrix, the off-diagonal elements cancel out, leaving only the diagonal elements to consider. Since the eigenvalues of a diagonal matrix are simply its diagonal elements, we can conclude that if the excitatory and inhibitory subpopulations are independently contracting (*w* is less than the contraction rate of an isolated neuron), then overall contraction is guaranteed. It is straightforward to generalize this simple two-neuron example to circuits achieving E-I balance through interacting *populations* (see A.5). It is also straightforward to generalize to the case where E-I and I-E connections do not cancel out exactly neuron by neuron, but rather they cancel out in a statistical sense where the mean amplitudes are matched. Another way to view this is E-I balance is in the framework of combinations of contracting systems (Fig 4). It is known that combining independently contracting systems in negative feedback preserves contraction^14^. We show that E-I balance actually translates to this negative feedback and thus can preserve contraction.

**Figure 4:**
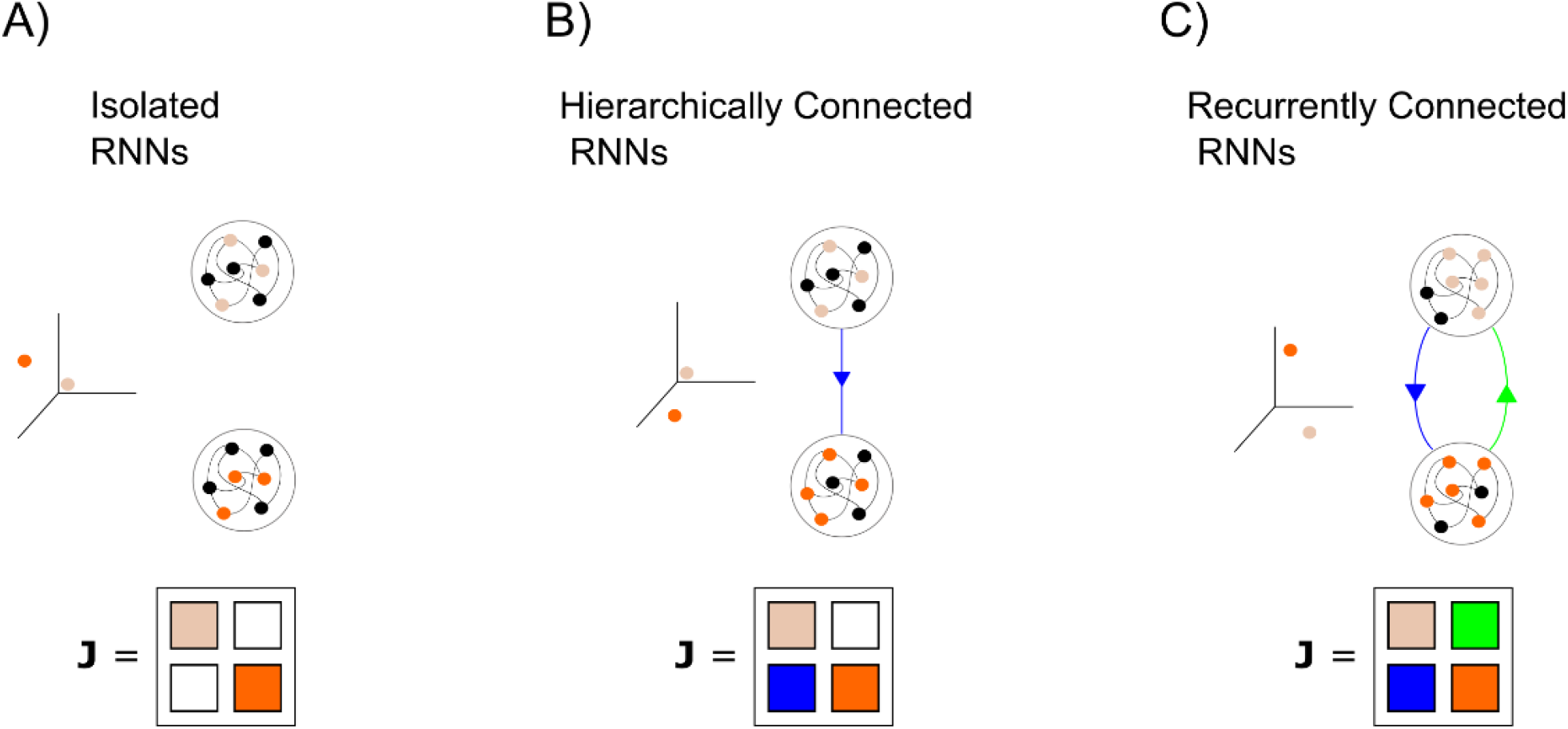
Combination properties of contracting systems. A) Two isolated, autonomous networks. The Jacobian of the overall system is block diagonal B) If one of the systems is connected to the other in a feedforward manner, the fixed point of the ‘bottom

### 3.4 Echo-State Networks Are Special Cases of Contracting RNNs

As can be seen in Figure 2, contracting systems have ‘fading memories’. This means that past events will affect the current state, but that the impact of a transient perturbation gradually decays over time. Consider the transient input in Figure 2 (red panel) presented on only one of the two trials to the network. Because the input is only present on one trial and not the other we call it a perturbation. When this perturbation occurs, the trajectories of the two trials become separated. However, after the disturbance is removed, the distance between the network’s trajectories starts shrinking back to zero again.

Thus, the network does not hold onto the memory of the perturbation indefinitely—the memory fades away. A similar property has been used in Echo State Networks (ESNs) to perform useful brain-inspired computations^38,39^. These networks are an alternative to classical attractor models in which neural computations are performed by entering stable states rather than by ‘fading memories’ of external inputs^40^. Because of the ‘fading memory’ property displayed by our contracting systems, we suspected that they might be related to ESNs. We investigated this next.

There are several distinctions between the networks described here and ESNs: 1) ESNs are discrete-time dynamical systems. This means that their states do not evolve continuously with time, but rather in ‘steps’. We primarily consider continuous time networks here. While attempts have been made to find ‘Echo-State Properties’ for leaky-integrator RNNs, these have all have relied on discretization of the continuous dynamics^41^. 2) To our knowledge, there are no known networks with dynamic synapses which provably have the ESP and 3) of the networks which are known to have ESP, they are all contracting in a static metric. This means that the “yardstick” by which distances are measured in an ESNs state space never change, thus limiting the scope of networks classifiable as ESNs. By removing dynamic synapses, setting the metric we use to prove contraction equal to the identity metric, and switching to a discrete time RNN, we could derive the so-called ‘Echo state condition’ as a special case of the contracting networks considered here (see A.5.1). It therefore follows that all the useful neural computations that have been performed by ESNs can automatically be performed by special instances of the networks considered in our work. However, by working within the framework of contraction analysis we were able to study networks both with dynamic synapses and non-identity metrics. This allowed for greater complexity in the network dynamics while preserving the “fading memory” property.

## 4 Discussion

We studied a fundamental question in neuroscience: How do neural circuits maintain stable dynamics in the presence of disturbance, noisy inputs and plastic change? We approached this problem from the perspective of dynamical systems theory, in light of the recent successes of understand neural circuits as dynamical systems^42^. We focused on *contracting* dynamical systems, which are yet largely unexplored in neuroscience, as a solution to the problem outlined above. We did so for three reasons:

1. Contracting systems can be *input-driven*. This is important because neural circuits are typically bombarded with time-varying inputs either from the environment or from other brain areas. Previous stability analyses have focused primarily on the stability of RNNs without time-varying input. These analyses are most insightful in situations where the input into a circuit can be approximated as either absent or constant. However, naturalistic stimuli tend to be highly time-varying and complex^43^.
2. Contracting systems are robust to noise and disturbances. Perturbations to a contracting system are forgotten at the rate of the contraction and noise therefore does not stack up over time. Thus dynamic stability can co-exist with high trial-to-trial variability in contracting neural networks, as observed in biology.
3. Contracting systems can be combined with one another in ways that preserve contraction (Fig 4). This is not true of most dynamical systems which can easily ‘blow up’ when connected in feedback with one another^8^. This combination property is important as it is increasingly clear that cognitive functions such as working memory or attention are distributed in multiple cortical and sub-cortical regions^44,45^. In particular, prefrontal cortex has been suggested as a hub that can reconfigure the cortical effective network based on task demands^46^. Brain networks must therefore be able to effectively reconfigure themselves on a fast time-scale without loss of stability. Most attempts in modelling cognition, for instance working memory, tend to utilize single and often autonomous networks. Contracting networks display a combination of input-driven and autonomous dynamics, and thus have key features necessary for combining modules into flexible and distributed networks.

To understand what mechanisms lead to contraction in neural circuits, we applied contraction analysis to RNNs. For RNNs with static weights, we found that the well-known Echo State Networks are a special case of a contracting network. Since realistic synapses are complex dynamical systems in their own right, we went one step further and asked when neural circuits with dynamic synapses would be contracting. We found that inhibitory Hebbian plasticity as well as excitatory anti-Hebbian plasticity and synaptic sparsity all lead to contraction in a broad class of RNNs.

Inhibitory plasticity has recently been the focus of many experimental and computational studies due to its stabilizing nature as well as its capacity for facilitating nontrivial computations in neural circuits^27,28,47^. It is known to give rise to excitatory-inhibitory balance and has been implicated as the mechanism behind many experimental findings such as sparse firing rates in cortex^28^. Similarly, anti-Hebbian plasticity exists across many brain areas and species, such as salamander and rabbit retina^29^, rat hippocampus^48,49^ electric fish electrosensory lobe^50^ and mouse prefrontal cortex^51^. Anti-hebbian dynamics can give rise to sparse neural codes which decrease correlations between neural activity and increase overall stimulus representation in the network^52^. Because of this on-line decorrelation property, anti-Hebbian plasticity has also been implicated in predictive coding^29,50^. Our findings suggest that it also increase the stability of networks.

For more general forms of synaptic dynamics, we showed (in section 3.2) that synaptic sparsity pushes RNNs towards being contracting. This aligns well with the experimental observation that synaptic connectivity is typically extremely sparse in the brain. Our results suggest that sparsity may be one factor pushing the brain towards dynamical stability. It is therefore interesting that synapses are regulated by homeostatic processes where synapses neighboring an upregulated synapse are immediately downregulated^53^. On the same note, we also observed that balancing the connections between excitatory and inhibitory populations leads to contraction. Balance between excitatory and inhibitory synaptic inputs are often observed in biology^34–36^ and could thus serve contractive stability purposes. Related computational work on spiking networks has suggested that balanced synaptic currents leads to fast response properties, efficient coding, increased robustness of function and can support complex dynamics related to movements^21,54–56^.

Experimental neuroscience is moving in the direction of studying many interacting neural circuits simultaneously. This is fueled by the expanding capabilities of recording multiple areas simultaneously in vivo and study their interactions. This increase the need for multi-modal cognitive models. We therefore anticipate that the presented work can provide a useful foundation for how cognition in noisy and distributed computational networks can be understood.

## Materials and Methods

All detailed proofs of main results are found in the appendix. Simulations (Figures 2 and 3) were performed in Python. Code to reproduce the figures is available at [https://github.com/kozleo/stable_dynamics]. Numerical integration was performed using sdeint, an open-source collection of numerical algorithms for integrations stochastic ordinary differential equations.

### Figure 2 details

All parameters and time constants in equations (1) and (2) were set to one. The integration step-size, *dt*, was set to 1e-2.

Initial conditions for both neural and synaptic activation were drawn uniformly between −1 and 1. Inputs into the network were generated by drawing *N* frequencies uniformly between *dt* and 100*dt*, phases between 0 and 2*π*, amplitudes between 0 and 20 and generating an N x *Time* vector of sinusoids with the above parameters.

The perturbations of the network was achieved by adding a vector of all 10s (i.e an additive vector input into the network, with each network of the element equal to 10) to the above input on one of the trials for 100 time steps in the middle of the simulation.

The noise was generated by driving each neural unit with an independent Weiner process (sigma = .2).

### Figure 3 details

The weight matrix used was the same as in Figure 2, leftmost panel (without perturbation, without noise).

## Supporting Information Legend

The supplementary appendix file contains extensive mathematical proofs of the results stated above. We kept the appendix self-contained by restating the basic results of contraction analysis and linear algebra which we used often in our proofs

## Acknowledgments

We thank Pawel Herman for comments on an earlier version of this manuscript. We thank Michael Happ and all members of the Miller Lab for helpful discussions and suggestions. This work was supported by NIMH R37MH087027, ONR MURI N00014-16-1-2832, NSF 1809314, and The MIT Picower Institute Innovation Fund.

## References

1. Lundqvist, M. et al. Gamma and Beta Bursts Underlie Working Memory. Neuron 90, 152–164 (2016).

2. Churchland, M. M. et al. Stimulus onset quenches neural variability: A widespread cortical phenomenon. Nat. Neurosci. 13, 369–378 (2010).

3. Hopfield, J. J. Neural networks and physical systems with emergent collective computational abilities. Proc. Natl. Acad. Sci. 79, 2554–2558 (1982).

4. Hirsch, M. W. Convergent activation dynamics in continuous time networks. Neural Networks 2, 331–349 (1989).

5. Cohen, M. A. & Grossberg, S. Absolute Stability of Global Pattern Formation and Parallel Memory Storage by Competitive Neural Networks. IEEE Trans. Syst. Man Cybern. SMC-13, 815–826 (1983).

6. Lundqvist, M., Herman, P. & Lansner, A. Theta and gamma power increases and alpha/beta power decreases with memory load in an attractor network model. J. Cogn. Neurosci. 23, 3008–3020 (2011).

7. Lansner, A. & Ekeberg, O. Reliability and Speed of Recall in an Associative Network. IEEE Trans. Pattern Anal. Mach. Intell. PAMI-7, 490–498 (1985).

8. Ashby, W. Design for a brain: The origin of adaptive behaviour. (Chapman & Hall Ltd, 1952).

9. Dayan, P. & Abbot, L. F. Theoretical Neuroscience Computational Neuroscience. The MIT press 241, (2005).

10. Zhang, H., Wang, Z. & Liu, D. A comprehensive review of stability analysis of continuous-time recurrent neural networks. IEEE Trans. Neural Networks Learn. Syst. 25, 1229–1262 (2014).

11. Spaak, E., Watanabe, K., Funahashi, S. & Stokes, M. G. Stable and dynamic coding for working memory in primate prefrontal cortex. J. Neurosci. 37, 6503–6516 (2017).

12. Laje, R. & Buonomano, D. V. Robust timing and motor patterns by taming chaos in recurrent neural networks. Nat. Neurosci. 16, 925–933 (2013).

13. Chaisangmongkon, W., Swaminathan, S. K., Freedman, D. J. & Wang, X. J. Computing by Robust Transience: How the Fronto-Parietal Network Performs Sequential, Category-Based Decisions. Neuron 93, 1504–1517.e4 (2017).

14. Lohmiller, W. & Slotine, J.-J. E. On Contraction Analysis for Non-linear Systems. Automatica 34, 683–696 (1998).

15. Rutishauser, U., Douglas, R. J. & Slotine, J.-J. Collective stability of networks of winner-take-all circuits*. (2018). doi:10.1162/NECO

16. Rutishauser, U., Slotine, J.-J. & Douglas, R. Computation in Dynamically Bounded Asymmetric Systems. PLoS Comput Biol 11, 1004039 (2015).

17. Girard, B., Tabareau, N., Pham, Q. C., Berthoz, A. & Slotine, J.-J. Where neuroscience and dynamic system theory meet autonomous robotics: A contracting basal ganglia model for action selection. Neural Networks 21, 628–641 (2008).

18. Tabareau, N., Slotine, J. J. & Pham, Q. C. How synchronization protects from noise. PLoS Comput. Biol. 6, 1–9 (2010).

19. Orhan, A. E. & Ma, W. J. A diverse range of factors affect the nature of neural representations underlying short-term memory. Nat. Neurosci. 22, 275–283 (2019).

20. Mongillo, G., Barak, O. & Tsodyks, M. Synaptic Theory of Working Memory. Science (80-.). 319, 1543 (2008).

21. Lundqvist, M., Compte, A. & Lansner, A. Bistable, Irregular Firing and Population Oscillations in a Modular Attractor Memory Network. PLoS Comput. Biol. 6, e1000803 (2010).

22. Söderlind, G. The logarithmic norm. History and modern theory. BIT Numer. Math. 46, 631–652 (2006).

23. Hennequin, G., Vogels, T. P. & Gerstner, W. Optimal control of transient dynamics in balanced networks supports generation of complex movements. Neuron 82, 1394–1406 (2014).

24. Stroud, J. P., Porter, M. A., Hennequin, G. & Vogels, T. P. Motor primitives in space and time via targeted gain modulation in cortical networks. Nat. Neurosci. 21, 1774–1783 (2018).

25. Zenke, F., Gerstner, W. & Ganguli, S. The temporal paradox of Hebbian learning and homeostatic plasticity. Current Opinion in Neurobiology 43, 166–176 (2017).

26. Vogelsy, T. P. et al. Inhibitory synaptic plasticity: Spike timing-dependence and putative network function. Frontiers in Neural Circuits (2013). doi:10.3389/fncir.2013.00119

27. Hennequin, G., Agnes, E. J. & Vogels, T. P. Inhibitory Plasticity: Balance, Control, and Codependence. Annu. Rev. Neurosci. 40, 557–579 (2017).

28. Vogels, T. P., Sprekeler, H., Zenke, F., Clopath, C. & Gerstner, W. Inhibitory Plasticity Balances Excitation and Inhibition in Sensory Pathways and Memory Networks. Science (80-.). 334, 1569–1573 (2011).

29. Hosoya, T., Baccus, S. A. & Meister, M. Dynamic predictive coding by the retina. Nature 436, 71 (2005).

30. Gerstner, W. & Kistler, W. M. Mathematical formulations of Hebbian learning. Biol. Cybern. 87, 404–415 (2002).

31. Slotine, J. J. E. Modular stability tools for distributed computation and control. Int. J. Adapt. Control Signal Process. 17, 397–416 (2003).

32. Kandel, E. R. et al. Principles of neural science. 4, (McGraw-hill New York, 2000).

33. Song, S., Sjöström, P. J., Reigl, M., Nelson, S. & Chklovskii, D. B. Highly nonrandom features of synaptic connectivity in local cortical circuits. PLoS Biol. 3, 0507–0519 (2005).

34. Mariño, J. et al. Invariant computations in local cortical networks with balanced excitation and inhibition. Nat. Neurosci. 8, 194–201 (2005).

35. Wehr, M. & Zador, A. M. Balanced inhibition underlies tuning and sharpens spike timing in auditory cortex. Nature 426, 442–446 (2003).

36. Shu, Y., Hasenstaub, A. & McCormick, D. A. Turning on and off recurrent balanced cortical activity. Nature 423, 288–293 (2003).

37. Murphy, B. K. & Miller, K. D. Balanced Amplification: A New Mechanism of Selective Amplification of Neural Activity Patterns. Neuron 61, 635–648 (2009).

38. Jaeger, H. The “echo state” approach to analysing and training recurrent neural networks-with an erratum note. Bonn, Ger. Ger. Natl. Res. Cent. Inf. Technol. GMD Tech. Rep. 148, 13 (2001).

39. Pascanu, R. & Jaeger, H. A Neurodynamical Model for Working Memory.

40. Buonomano, D. V & Maass, W. State-dependent computations: spatiotemporal processing in cortical networks. (2009). doi:10.1038/nrn2558

41. Jaeger, H., Lukoševičius, M., Popovici, D. & Siewert, U. Optimization and applications of echo state networks with leaky-integrator neurons. Neural Networks 20, 335–352 (2007).

42. Sussillo, D. Neural circuits as computational dynamical systems. Curr. Opin. Neurobiol. 25, 156–163 (2014).

43. Steveninck, R. R. D. R. Van, Lewen, G. D., Strong, S. P., Koberle, R. & Bialek, W. Reproducibility and Variability in Neural Spike Trains. 275, (1997).

44. Chatham, C. H. & Badre, D. Multiple gates on working memory. Curr. Opin. Behav. Sci. 1, 23–31 (2015).

45. Halassa, M. M. & Kastner, S. Thalamic functions in distributed cognitive control. Nat. Neurosci. 20, 1669–1679 (2017).

46. Miller, E. K. & Cohen, J. D. An Integrative Theory of Prefrontal Cortex Function. Annu. Rev. Neurosci. 24, 167–202 (2001).

47. Vogelsy, T. P. et al. Inhibitory synaptic plasticity: Spike timing-dependence and putative network function. Front. Neural Circuits 7, 1–11 (2013).

48. Lisman, J. A mechanism for the Hebb and the anti-Hebb processes underlying learning and memory. Proc. Natl. Acad. Sci. 86, 9574–9578 (1989).

49. Kullmann, D. M. & Lamsa, K. P. Long-term synaptic plasticity in hippocampal interneurons. Nat. Rev. Neurosci. 8, 687–699 (2007).

50. Enikolopov, A. G., Abbott, L. & Sawtell, N. B. Internally Generated Predictions Enhance Neural and Behavioral Detection of Sensory Stimuli in an Electric Fish. (2018). doi:10.1016/j.neuron.2018.06.006

51. Ruan, H., Saur, T. & Yao, W.-D. Dopamine-enabled anti-Hebbian timing-dependent plasticity in prefrontal circuitry. Front. Neural Circuits 8, 38 (2014).

52. Földiák, P. Forming sparse representations by local anti-Hebbian learning. Biol. Cybern. 64, 165–170 (1990).

53. El-Boustani, S. et al. Locally coordinated synaptic plasticity of visual cortex neurons in vivo. Science (80-.). 360, 1349–1354 (2018).

54. Denève, S. & Machens, C. K. Efficient codes and balanced networks. Nat. Neurosci. 19, 375–382 (2016).

55. Hennequin, G., Vogels, T. P. & Gerstner, W. Optimal control of transient dynamics in balanced networks supports generation of complex movements. Neuron 82, 1394–1406 (2014).

56. Brunel, N. Dynamics of Sparsely Connected Networks of Excitatory and Inhibitory Spiking Neurons. Journal of Computational Neuroscience 8, (2000).

